# Anaerobic single particle cryoEM of nitrogenase

**DOI:** 10.1101/2022.06.04.494841

**Authors:** Rebeccah A. Warmack, Douglas C. Rees

**Affiliations:** Division of Chemistry and Chemical Engineering 147-75, California Institute of Technology, Pasadena, California 91125, United States; Howard Hughes Medical Institute, California Institute of Technology, United States

## Abstract

The enzyme nitrogenase catalyzes the reduction of dinitrogen to ammonia during biological nitrogen fixation through a mechanism involving the ATP dependent interaction of two component proteins adopting multiple conformational states. To date, high resolution structural information has been provided by X-ray crystallography, which restricts the states that can be accessed to those that can be crystallized. Cryo-electron microscopy (cryoEM) presents a new opportunity for structural characterization of nitrogenase solution structures, and may yield new information on the mechanism of nitrogenase by revealing structures of transient or heterogeneous states. In this study, we present single particle cryoEM structures of the MoFe-nitrogenase endogenously isolated from *Azotobacter vinelandii*. To maintain the fully reduced cluster states of this oxygen sensitive protein, we prepared samples within an anaerobic chamber and employed specialized conditions to minimize partial disordering of the α-subunit at the air-water interface during freezing. Under these conditions, cryoEM structures of the as-isolated MoFe-protein and stabilized MoFe-protein-Fe-protein ADP-AlF_4_^-^complex were generally found to closely resemble their corresponding X-ray crystallographic structures. The cryoEM structures did reveal disordering in regions of the MoFe-protein α-subunit reminiscent of that observed previously for the Δ*nifB* MoFe-protein lacking the FeMo-cofactor, suggesting that this disorder may reflect functionally relevant dynamics, as well as the possibility of asymmetric binding of the Fe-protein to the MoFe-protein in solution. The methods presented here pave the way toward the capture and interrogation of turnover-relevant nitrogenase states by cryoEM.

## Introduction

A thorough understanding of the catalytic strategy employed by biological nitrogen fixation (BNF) to reduce dinitrogen to ammonia under ambient conditions may inform the development of more efficient ammonia production platforms. BNF occurs within diazotrophic organisms primarily via the molybdenum iron (MoFe) nitrogenase (reviewed in (*1-5*)), though other homologs, including the vanadium and iron-only nitrogenases, are expressed under Mo-limiting conditions (*6,7*). The MoFe-nitrogenase consists of two proteins, the molybdenum iron (MoFe-) protein, containing the active site metallocluster for dinitrogen reduction, and the iron (Fe-) protein that mediates the ATP-dependent transfer of electrons to the MoFe-protein. The MoFe-protein is a 230 kDa α_2_β_2_ heterotetramer with two complex metalloclusters per αβ-dimer: the [8Fe:7S] P-cluster positioned between the α- and β-subunits, and the [7Fe:1Mo:9S:1C]-*R*-homocitrate FeMo-cofactor coordinated within the α-subunit. The Fe-protein is a 64 kDa γ_2_ homodimeric nucleotide-binding protein containing a [4Fe:4S] cluster required for electron transfer to the MoFe-protein. Kinetic studies have established that the coupling of ATP hydrolysis to electron transfer occurs in the MoFe-protein - Fe-protein complex, with a stoichiometry conventionally represented as two ATP hydrolyzed per electron transferred. Electrons are subsequently funneled into the active site cluster of the MoFe-protein, the FeMo-cofactor (*8*). Substrate reduction requires multiple cycles of this Fe-protein cycle, involving repeated association, electron-transfer, dissociation, and ATP-reloading of the Fe-protein (*8,9*); significantly, the Fe-protein is the only known reductant capable of efficiently supporting N_2_ reduction at the MoFe-protein active site.

Structures of the as-isolated states of the MoFe-protein from *Azotobacter vinelandii* alone and in complex with the Fe-protein have been solved to high resolution by X-ray crystallography (*10-14*). Subsequently determined structures of inhibitor-bound or selenium-incorporated MoFe-protein have demonstrated the labile nature of belt sulfurs within the FeMo-cofactor (*15-18*). However, the visualization of transiently occurring states that are critical to defining the catalytic mechanism remains elusive, reflecting the dynamic nature of the nitrogenase system under turnover conditions. Single particle cryoEM represents a new avenue for the structural investigation of nitrogenase, as it has become an increasingly high resolution technique (*19-20*). This method requires very low protein concentrations and the sample preparation process requires rapid vitrification of protein in solution that may better trap intermediate states than is possible by crystallization. These characteristics make single particle cryoEM an ideal method to explore transient intermediate, reduced, or substrate bound states of nitrogenase. However, the applicability of this method to the extremely oxygen sensitive nitrogenase proteins (*21*) has not yet been demonstrated.

We report the development of methods to reconstruct single particle cryo-EM structures of the intact, as-isolated reduced state of the MoFe-protein. These methods included placing a plunge freezing robot in an anaerobic chamber to maintain the fully reduced cluster states of this oxygen sensitive protein, and the use of EM grids with ultrathin carbon or detergent to minimize disordering of the a-subunit at the air-water interface during freezing. Interestingly, reconstructions of the AWI-perturbed MoFe-protein reveal α-subunit disordering that resembles that observed previously for the Δ*nifB* MoFe-protein lacking the FeMo-cofactor (*22*), suggesting that this disorder may reflect functionally relevant dynamics. These conditions consistently yield sub 3 Å resolution structures of the fully reduced solution state MoFe-protein alone and in the ADP-AlF_4_^-^stabilized complex with the Fe-protein, illustrating the applicability of single particle cryoEM for the structural study of nitrogenase catalysis.

## Results

### Disorder is induced in *A. vinelandii* MoFe-nitrogenase by the air-water interface on EM grids

Our initial attempts to determine the structure of the *Azotobacter vinelandii* MoFe-protein by single particle cryo-EM utilized holey carbon grids prepared on a Vitrobot housed within an anaerobic chamber. The micrographs showed a dense, homogenous population of the MoFe-protein, and yielded a 2.56 Å resolution map showing asymmetrically disordered regions (Figure 1A; Supp. Figure 1A). Comparison of this volume to a previous crystal structure of MoFe-protein established a region of missing electrostatic potential (ESP) largely within the a-subunit primarily within one a dimer, which we have termed the “disordered” dimer (Figure 1A). This partial disordering is reminiscent of that observed in the structure of the cofactor-deficient Δ*nifB* MoFe-protein (Figure 1B; ref. *22*, PDB code 1L5H). Indeed, rigid body fitting of this structure into the EM ESP showed significant overlap between regions of missing ESP and the disordered residues within this FeMo-cofactor deficient structure (Figure 1B). Specifically, the EM map lacks ESP in the region corresponding to residues 1-52 and 375-416 of domain III in the α-subunit, closely corresponding to the disorder at residues 1-48, 381-407, and 481-492 in the Δ*nif* species (Figure 1C-D). Intriguingly, while the FeMo-cofactor ESP on the “ordered” dimer displays the canonical geometry and overlaps with the crystallographic location of the cluster in the holo protein, the ESP corresponding to the FeMo-cofactor on the “disordered” dimer is significantly shifted towards the surface of the protein (Figure 1E). Local resolution indicates low resolution around this cluster region and at higher thresholds the ESP for this FeMo-cofactor disappears rapidly, suggestive of positional disorder at this site.

**Figure 1:**
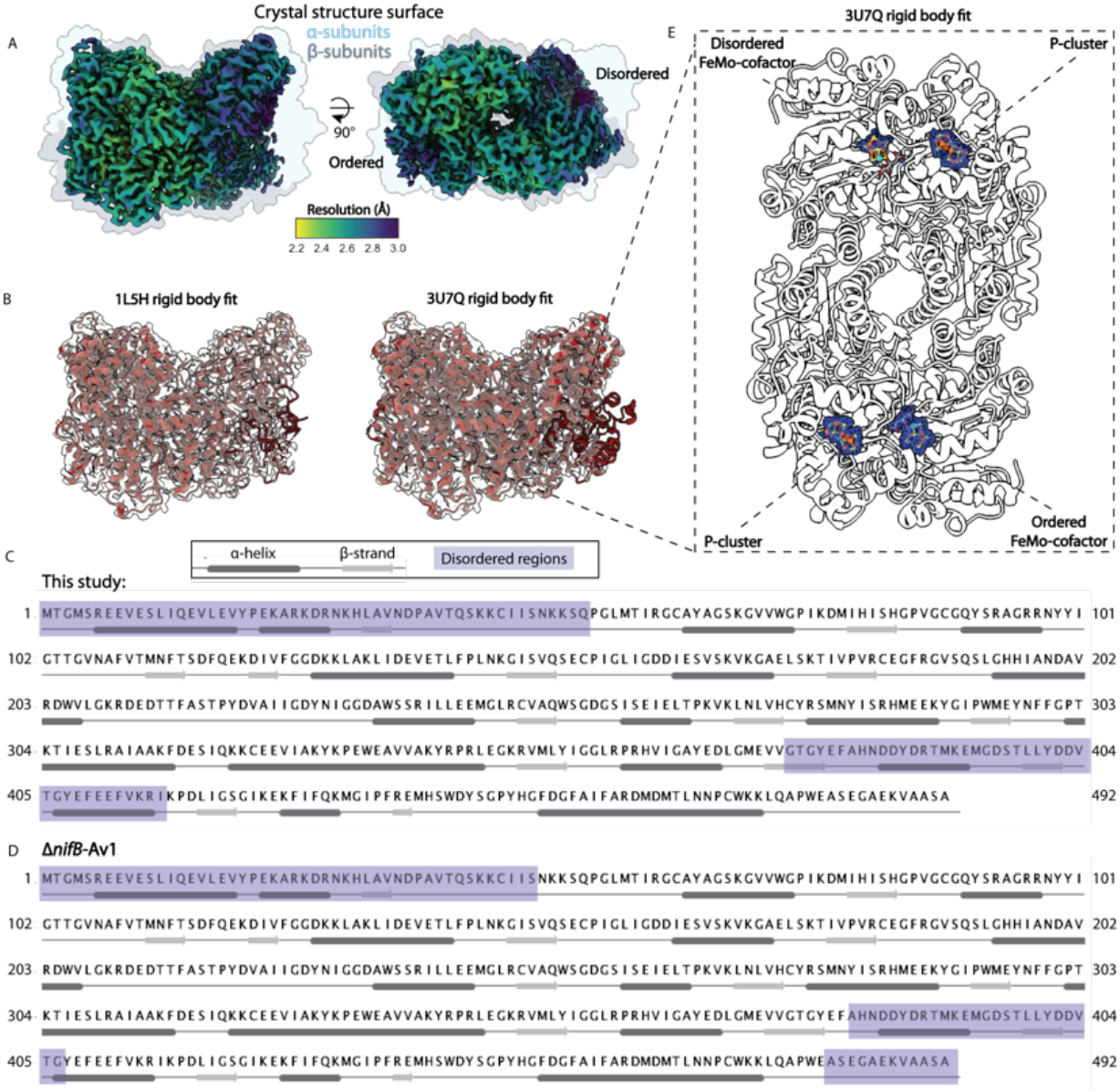
Air-water interface damaged cryoEM map of the MoFe-protein. (a) 2.56 Å resolution (C1) cryoEM map of the MoFe-protein shaded according to local resolution overlaid with surface representation of MoFe-protein crystal structure (PDB code 3U7Q). (b) Rigid body fit of the as-isolated MoFe-protein crystal structure (PDB code 3U7Q) and the FeMo-cofactor deficient MoFe-protein crystal structure (Δ*nifB*-Av1; PDB code 1L5H) in the cryoEM ESP. (c-d) Sequence of the MoFe-protein alpha subunit with disordered residues highlighted from the cryoEM map and the Δ*nifB*-Av1 protein, respectively. Secondary structure was annotated with jpred (ref. 23). (e) CryoEM ESP of the metalloclusters overlaid with a rigid body fit of PDB code 3U7Q.

Examination of the single particle grids by cryogenic electron-tomography (cryoET) revealed that in regions of thin ice (∼20 nm) particles form a monolayer in which most particles lie within 5-10 nm of the air-water interface (AWI) (Supp. Fig 1C). In thicker ice layers (>100 nm), MoFe-protein particles still localize to the AWI on either face of the vitreous ice, with few particles distributed within the ice layer (Figure 2A-B). Analysis of the 2D classes from these conditions during reconstruction reveal loss of features corresponding to the α-subunit as seen in the full reconstruction (Figure 2B, lower panel). Significantly, the particle orientations apparent in the final subset reflect strong preferred orientation corresponding to the disordered region, suggesting that the α-subunit is oriented towards the AWI (Figure 2C).

**Figure 2:**
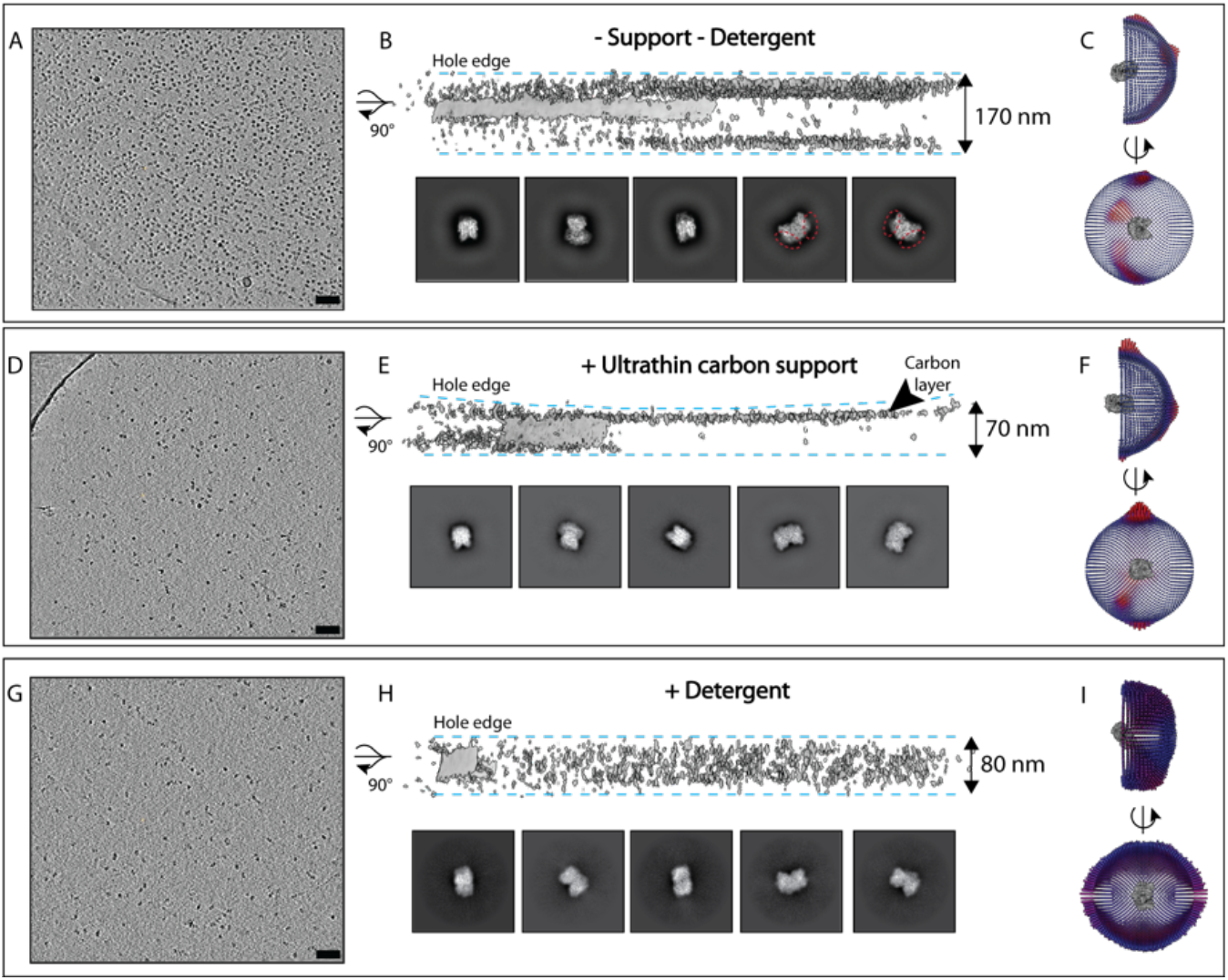
Detergent supplementation limits air water interface adsorption and preferred orientation of MoFe-nitrogenase particles. Top panel: MoFe-protein particles alone on holey carbon grids. Middle panel: MoFe-protein particles alone on ultrathin carbon layered grids. Bottom panel: MoFe-protein particles with detergent on holey carbon grids. (a, d, g) reconstructed tomogram of MoFe-protein particles. (b, e, h) Top: Volumetric representation of tomogram showing particle distribution in ice, scale bars represent 100 nm. Middle panel, bottom: Representative 2D classes from corresponding single particle reconstructions. (c, f, i) Euler angle distributions from reconstructed single particle cryoEM maps (Red, overrepresented views).

As adsorption of proteins to the AWI has been associated with protein denaturation (*24*), we screened various methods of grid preparation to mitigate this behavior. Both carbon-layered grids and detergent-supplemented conditions were found to minimize the adsorption of particles to the AWI. In the case of carbon-layered grids, MoFe-protein particles appear to preferentially adsorb to the carbon layer during freezing, thereby protecting the particles from interactions at the air water interface (Figure 2D-E). The particles lay on the carbon in a preferred orientation with the long axis of the particle lying against the grid (Figure 2F). Particles were more evenly distributed throughout the ice in detergent-containing conditions, perhaps supporting the formation of a protective detergent monolayer at the AWI (Figure 2G-H). The more nearly random distribution of the particles in the detergent condition is reflected in the large variety of views observed in the particle orientations (Figure 2I). Either detergent-containing conditions or carbon-layered grids were used in all subsequent experiments to circumvent structural artifacts that might result from AWI-interactions.

### As-isolated single particle cryoEM structure of MoFe-nitrogenase closely resembles its resting state X-ray crystal form

CryoEM grids prepared with the zwitterionic detergent CHAPSO yielded single particle cryoEM reconstructions of as-isolated MoFe-protein at 2.04 Å resolution, with higher resolution estimated within the metallocluster sites (Figure 3; Supp. Figure 2). An as-isolated MoFe-protein reconstruction at 2.26 Å resolution on carbon-layered grids was also obtained (Supp. Figure 3). Due to some residual preferred orientation of the carbon-adhered particles, the latter map displays slight anisotropy. The single particle cryoEM structure in CHAPSO revealed an all atom RMSD of 0.41 Å when compared to the 1 Å resolution X-ray structure from the Einsle group (PDB code 3U7Q), demonstrating that the overall conformation of the MoFe-protein polypeptide in crystals and in solution is similar. In accordance with these parallels, both the P-cluster and the FeMo-cofactor in the cryoEM structure align well with the 3U7Q model. The ESP for the P-cluster reveals the [8Fe:7S] bridged double cubane structure with a shared central sulfide assigned to the fully reduced P^N^ state, coordinated by residues Cys62, Cys88, and Cys154 in the α-subunit and residues Cys70, Cys95, and Cys153 in the β- subunit (Figure 3D). The FeMo-cofactor ESP within the α-subunit likewise exhibits the characteristic [7Fe:9S:Mo] structure organized around a trigonal prismatic arrangement of the central six irons, bridged by three belt sulfides. The inorganic core of the cluster is coordinated to the MoFe-protein α-subunit by Cys275 and His442, with the coordination sphere of the Mo completed by a bidentate homocitrate moiety (Figure 3D). While there is no ESP in the cryoEM map for the central carbide of the FeMo-cofactor, presumably due to resolution dependent effects, the conservation of the overall trigonal prism architecture implies the presence of this central ligand. In X-ray crystallography, this ligand was not observed until resolutions of 1.16 Å or higher were achieved due to Fourier series termination effects (*25*). The mononuclear metal binding site typically occupied by iron or calcium within the X-ray structures was also identified within the cryoEM map (Supp. Figure 2). Successive blurring of the map reveals that the cryoEM ESP around this site increases while surrounding atoms remain mostly constant, supporting the presence of a cation (*26*).

**Figure 3:**
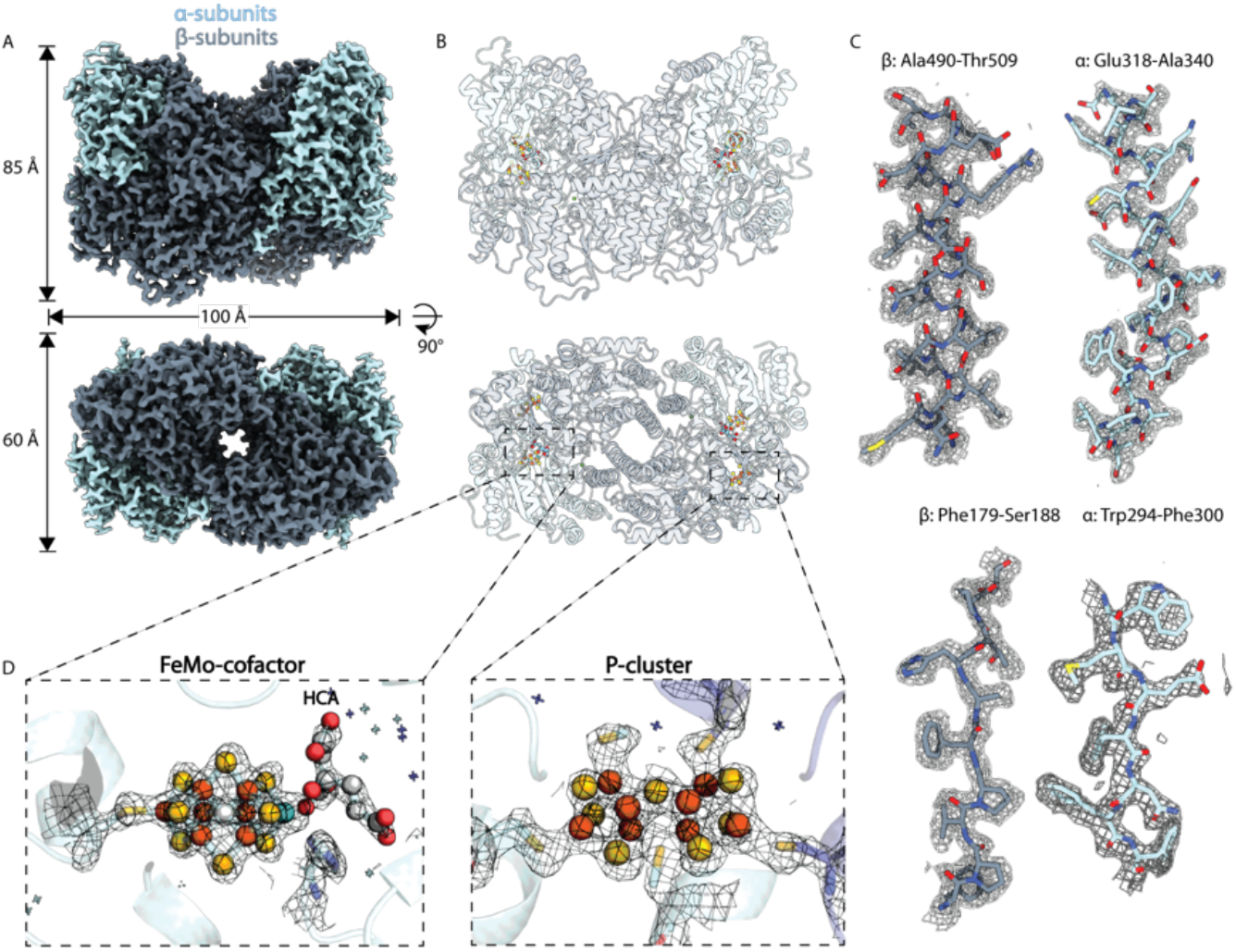
As-isolated MoFe-nitrogenase cryoEM structure in CHAPSO detergent. (a) CryoEM map of the anaerobically prepared MoFe-protein with detergent on holey carbon grids. (b) Model of MoFe-protein cryoEM structure. (c) Representative cryoEM ESP of secondary structure elements within the MoFe-protein α- and β-subunits. (d) CryoEM ESP around the modeled active site FeMo-cofactor and the P-cluster.

Portions of the α-subunit within the cryoEM map still display noticeably lower estimated local resolution in certain regions, along with higher B-factors for the corresponding residues in the structural model (Supp. Figure 2C). Within this detergent condition the effect is most noticeable at the N-terminus, with residues Asp36-Cys45 displaying particularly poor ESP. These low estimates of local resolution and high B-factors within the cryoEM structures may reflect a higher degree of flexibility or motion in the solution state α-subunit. The overlap in the region of partially disordered residues of the MoFe-protein within this detergent containing condition with the AWI-perturbed map and the Δ*nifB* MoFe-protein is notable.

Ordered waters in the protein interior of the cryoEM structure were found to overlap strongly with the X-ray structure. As waters have been suggested to participate in transport pathways within the MoFe-protein, waters surrounding the FeMo-cofactor are of particular interest. All of the waters identified within a 10 Å sphere of the cofactor in the solution structure are conserved in the X-ray structure. Relative to the X-ray structure, however, fewer ordered waters were identified on the surface of the cryoEM structure, perhaps reflecting the absence of lattice contacts in the latter that may contribute to more highly ordered waters present on the protein surface in the X-ray structure.

Interestingly, in this detergent condition, two distinct CHAPSO binding sites were observed in each αβ dimer (Supp. Figure 2). In both cases, the tail of the detergent is not resolved. One site rests between α-helix β39-48 and the loop extending from α142-147 (site 1), while the second site lies between α-helix α318-345, β-strand β414-416, loop β361-366, and loop β3 −11 (site 2). The backbones of these sites are unperturbed by the presence of the CHAPSO molecules in comparison to 3U7Q.

### Anaerobic conditions preserve the as-isolated reduced state

The process of preparing grids for cryoEM with commercial plunge freezers requires timescales on the order of several seconds. To preclude exposure to oxygen during this period, we placed a Vitrobot in an anaerobic chamber under an argon atmosphere. However, as working within an anaerobic chamber is cumbersome, to test whether these precautions are necessary, a sample of the MoFe-protein alone was transferred from a sealed anaerobic vial onto a grid within a benchtop Vitrobot (exposed to air) using a gastight syringe and immediately blotted and plunge frozen. Single particle cryoEM data collection and analysis of this grid yielded a 1.92 Å resolution map (Figure 4A). The model displayed an all atom RMSD of 0.65 Å with the high resolution crystal structure, and 0.48 Å with the anaerobic cryoEM structure (with detergent). Analysis of the ESP surrounding the P-cluster revealed bridging ESP between Fe6 and Ser β188 (Figure 4D, red asterisk). In the anaerobically purified as-isolated state of the MoFe protein, with the P-cluster in the reduced (P^N^) state, Fe6 and Fe5 are liganded to the sidechain thiols of Cys β153 and Cys α88, respectively. In the oxidized state of the MoFe-protein (P-cluster in the P^+2^ state), it has been shown that interactions with the central sulfur by Fe6 and Fe5 are replaced by the Ser β188 sidechain and the backbone amide of Cys α88, respectively (*27*). In 2018, Keable et al., crystallographically demonstrated that there is an intermediate P^+1^ state, in which the Fe6 becomes liganded to Ser β188, but the Fe5 remains coordinated to the central sulfur of the P-cluster (*28*). Consequently, the aerobically frozen MoFe-protein cryoEM structure corresponds to the oxidized P^+1^ state of the P-cluster, emphasizing the necessity for anaerobic cryoEM sample preparation conditions for the study of MoFe-nitrogenase.

**Figure 4:**
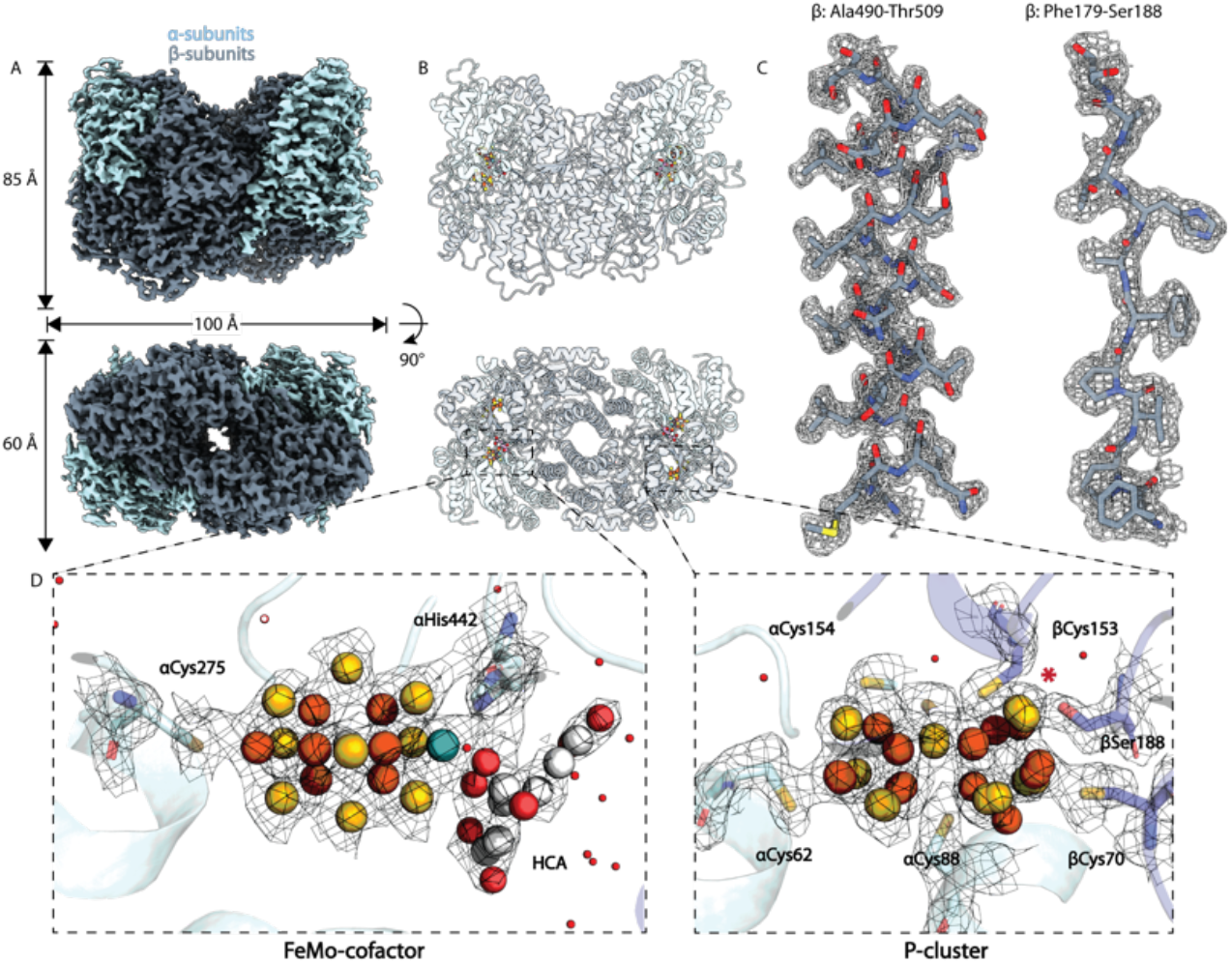
Partially oxidized MoFe-protein cryoEM structure. (a) CryoEM map of aerobically prepared MoFe-protein on carbon-layered grids. (b) Model of MoFe-protein cryoEM structure. (c) cryoEM ESP corresponding to residues within the MoFe-protein β-subunits. (d) Metallocluster cryoEM ESP and model. Red asterisk indicates bridging ESP between Ser β188 and Fe6.

Despite nominal ∼2 Å resolution, it should be noted that the local resolution of ESP within large regions of the α-subunit appear to be much lower, with the associated B-factors much higher than average (Supp. Figure 4). This disorder in combination with that observed within the as-isolated structure and the AWI-perturbed EM map further support of the possible native disorder within the α-subunit.

### Asymmetric, heterogeneous states observed in a non-resting state ADP-AlF_4_^-^stabilized MoFe-protein – Fe-protein complex

A key event in the nitrogenase mechanism is formation of a transient MoFe-protein – Fe-protein complex where ATP hydrolysis is coupled to electron transfer between the two proteins. This complex is inhibited by aluminum fluoride (AlF_4_^-^) with ADP, resulting in formation of a stable MoFe-protein – Fe-protein species that has been isolated and crystallographically characterized (*12-13*). To evaluate the applicability of cryoEM for the full nitrogenase complex, a reaction of the Fe-protein, MoFe-protein, ADP, AlCl_3_ and NaF was carried out as described in the Methods. Stabilized complex was isolated from this reaction mixture via size exclusion chromatography (SEC). Grids were prepared from the isolated ADP-AlF_4_^-^inhibited MoFe-protein – Fe-protein complex peak (Figure 5A). Interestingly, despite the isolation via SEC, replicate experiments and datasets, three distinct nitrogenase species were consistently observed on the grids with Fe-protein:MoFe-protein bound states at ratios of 2:1, 1:1, and 0:1 (Figure 5B). Classification of these separate states resulted in maps of the 2:1 and 1:1 Fe-protein:MoFe-protein complexes at 2.12 Å and 2.60 Å resolution, respectively (Figure 5C-D; Supp. Figure 5).

**Figure 5:**
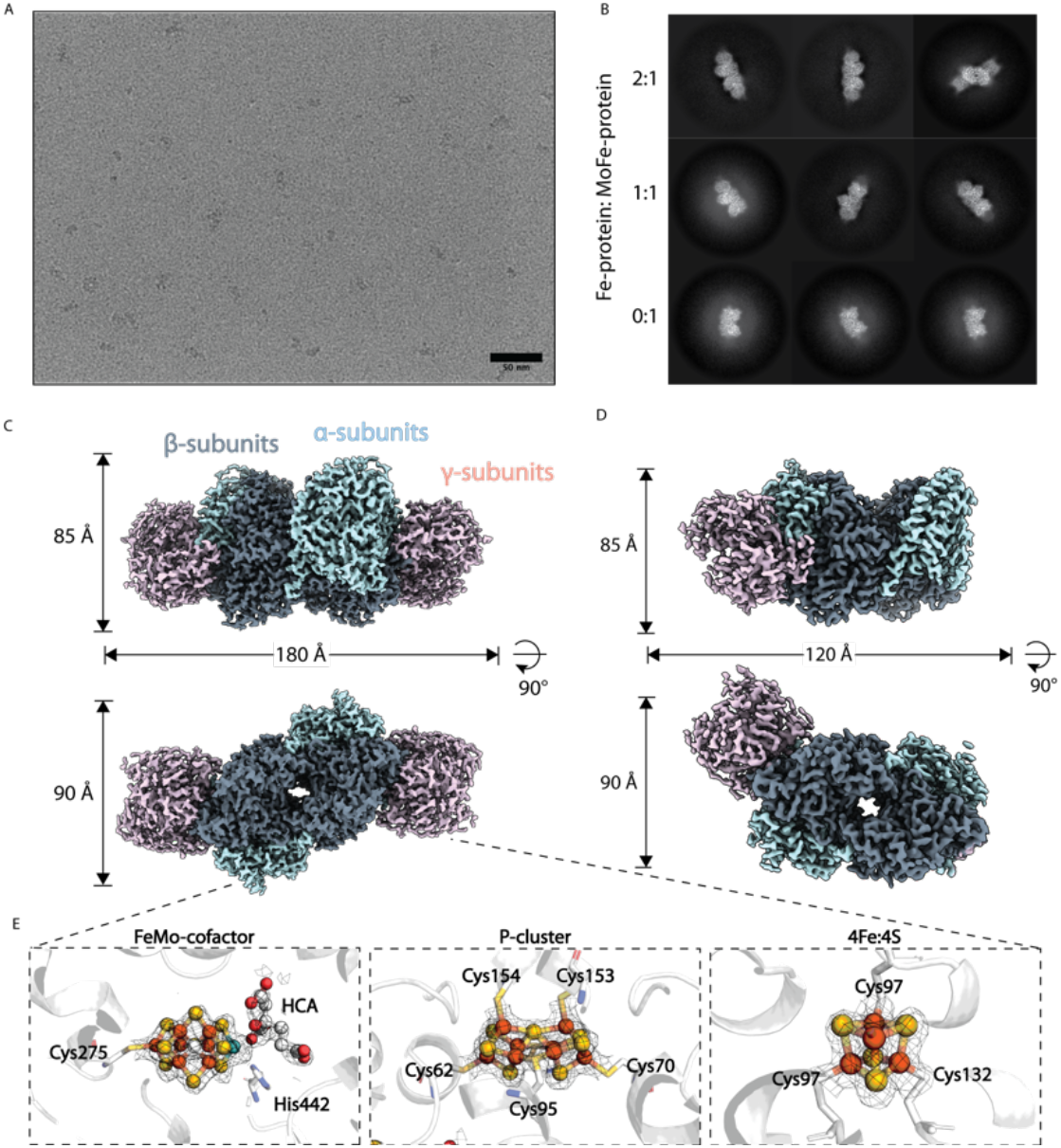
CryoEM structure of ADP-AlF_4_ ^-^ stabilized MoFe-protein-Fe-protein complex. (a) Representative micrograph of complex particles on ultrathin carbon layered grids. (b) 2D classes of three nitrogenase complex species with ratios of Fe-protein to MoFe-protein of 2:1, 1:1, and 0:1 (c) CryoEM map and model of the 2:1 ADP-AlF_4_^**-**^ stabilized MoFe-protein-Fe-protein complex. (d) CryoEM map and model of the 1:1 ADP-AlF_4_^**-**^ stabilized MoFe-protein-Fe-protein complex. (e) CryoEM ESP around the metalloclusters in the 2:1 complex.

As in the solution state cryoEM structure of the MoFe-protein alone, the 2:1 ADP-AlF_4_^-^MoFe-protein – Fe-protein complex maintains the two-fold axis relating the two αβ/αβγ halves, originally seen as a non-crystallographic two-fold axis in the crystal structures (Figure 5C). Similarly, in both the 2:1 and 1:1 complexes, residues in the N-termini and domain III of the MoFe-protein α-subunit also display poorer estimated local resolution within the maps and higher B-factors within the model. This disorder is more pronounced in the asymmetrically Fe-protein bound α-subunits of the 1:1 complex, with poorer ESP observed within the unbound αβ dimer (Figure 5D). Within the Fe-protein of both structures, the 4Fe:4S cluster remains liganded by residues Cys97 and Cys132 on the γ-subunits positioned approximately 14 Å away from the P-cluster between the α- and β- subunits. Both the P-cluster and the FeMo-cofactor within the full complex display expected geometries that align closely with the crystallographic locations of the metalloclusters in the previously reported X-ray crystal structure (*12*).

## Discussion

Structural studies of catalytic nitrogenase intermediates have long been limited by the need for highly pure and stable protein states for X-ray crystallography. In this study, we have demonstrated single particle cryo-EM as a high resolution method for the study of the mechanism of highly oxygen sensitive nitrogenase proteins which requires only small amounts of protein in solution. Optimization of this methodology for the nitrogenase system will pave the way for studies of nitrogenase turnover complexes, thus facilitating increased understanding of the enzymatic mechanism. A primary obstacle in working with nitrogenase is its extreme oxygen sensitivity. To overcome this barrier, we have constructed an anaerobic system for grid preparation by placing a grid freezing robot (Vitrobot) within an anaerobic chamber. These conditions have yielded high resolution cryoEM structures of the as-isolated MoFe-protein and the ADP-AlF_4_^-^ MoFe-protein – Fe-protein complex. Overall, these structures exhibit highly similar architecture to previously reported crystal structures. The similarity between the pseudo-solution-state cryoEM structures and the crystal structures indicates that the observed resting state represents a structure that can be primed for catalysis in solution.

While testing our cryoEM methodology with the as-isolated MoFe-protein alone, we discovered that this protein is susceptible to damage at the air-water interface (AWI), which results in disordering of the α-subunit and perturbations within the active site. Curiously, the missing regions of ESP within the AWI-damaged map correspond closely with disordered residues within the crystal structure of a FeMo-cofactor-deficient MoFe-protein isolated from a strain of *A. vinelandii* lacking the FeMo-cofactor biosynthetic protein nifB (ref. 22; Figure 1). The disordering was proposed to be relevant to cluster insertion processes, analogous to the opening of active sites for metal insertion in certain copper-containing proteins (*29-30*). Our EM maps demonstrate that disorder within similar regions in the α-subunit above the active site FeMo-cofactor is perpetuated to a lesser degree even in the absence of interactions with the air water interface. This disorder can manifest asymmetrically, particularly within the air water interface damaged form and in 1:1 ADP-AlF_4_^-^stabilized complexes of the Fe-protein and MoFe-protein.

The consistent nature of the disorder and its localization within regions associated with cofactor insertion may support a functional role for dynamics in the α-subunit, possibly during substrate reduction or perhaps participating in an as yet uncharacterized repair mechanism for replacing damaged cofactors. Studies have shown the labile nature of the belt sulfurs within the FeMo-cofactor and have suggested these sites are displaced and subsequently replaced during the catalytic cycle (*15-17*). These changes within the active site may require conformational rearrangements within the α-subunit to accommodate displaced sulfur containing species or altered FeMo-cofactor. Asymmetry in the disorder observed within the MoFe-protein α-subunit may additionally point to cooperativity between the two αβ dimers. We note that previous kinetic and biophysical studies (*31, 32*) have highlighted a possible role for negative cooperativity and 1:1 complexes in the interaction between the MoFe-and Fe-proteins.

The asymmetry of the α-subunit disorder is strikingly paralleled by our results with the ADP-AlF_4_^-^MoFe-protein – Fe-protein complex, with which we observe both symmetric 2:1 and asymmetric 1:1 complexes within a grid (Figure 5). In the asymmetric 1:1 complex, the α-subunit distal to the bound Fe-protein exhibits lower resolution and fragmented ESP. While cryoET reveals particles adsorb to the carbon layer on ultrathin carbon layered grids, it is possible that disorder within this exposed α-subunit is due to interactions at the AWI prior to adherence to the carbon layer. This might indicate that the Fe-protein is able to ‘protect’ the MoFe-protein from damage at the AWI. It is also possible that this asymmetric disorder is relevant to the bound state of the complex, and that the Fe-protein helps to order the α-subunit. While further studies are required to understand this relationship, these observations of the effect of the Fe-protein on α-subunit ordering may have direct relevance to dynamic turnover states.

The resolutions achieved in this study allowed us to further visualize changes within the metalloclusters of the MoFe-protein. While investigating the requirement of anaerobic conditions for cryoEM studies of nitrogenase, we discovered a partially oxidized form of the P-cluster within the MoFe-protein (Figure 4). Within the approximately 2 Å resolution map, we were able to visualize strong bridging ESP between Ser β188 and Fe6, but not between the amide of Cys α88 and Fe5, indicative of the partially oxidized P^+1^ state of the P-cluster (*27*). These results not only highlight the importance of anaerobic conditions for the preparation of cryoEM samples, but also demonstrate the potential applicability of cryoEM for the unambiguous identification of changes within the metalloclusters of the MoFe-protein during turnover.

In this study, we have demonstrated the applicability of single particle cryoEM for the study of nitrogenase. Importantly, our experiments show that anaerobic conditions are required to maintain the as-isolated reduced state of nitrogenase and that the protein is susceptible to damaging modifications at the AWI. We present methodology to circumvent both issues for future application of single particle cryoEM to the structural investigation of nitrogenase turnover states.

## Materials & Methods

### Purification of MoFe-Nitrogenase

The MoFe-protein and the Fe-protein were purified from *Azotobacter vinelandii* under anaerobic conditions using a combination of Schlenk line techniques and anaerobic chambers with oxygen-scrubbed argon as described previously (*33*). Protein concentration was determined by absorbance at 410 nm using extinction coefficients of 76 and 9.4 mM^-1^cm^-1^ for the MoFe-protein and the Fe-protein, respectively. Concentration calculations were based on molecular weights of 230 and 64 kDa for the MoFe protein and the Fe-protein, respectively. The specific activities for acetylene reduction was ∼2300 nmol/min/mg MoFe-protein. Component ratio (CR) of Fe-protein to MoFe-protein was defined as moles of Fe-protein per mole of MoFe-protein active site via the equation CR=1.82(*C*_*Fe*_*/C*_*MoFe*_), where *C*_*Fe*_ and *C*_*MoFe*_ are the concentrations of the Fe-protein and MoFe-protein in the final reaction mixture, respectively, in mg/mL.

### Preparation of the stabilized complex

ADP-AlF_4_^-^stabilized Fe-protein-MoFe-protein complex was prepared as described in Renner and Howard (*34*). Briefly, 0.25 mg MoFe-protein and Fe-protein at a CR of ∼2 in a final volume of 1.3 mL were incubated in 100 mM MOPS, 50 mM Tris, 100 mM NaCl (pH 7.3) with 10 mM sodium dithionite, 4 mM NaF, 0.2 mM AlCl_3_, 1 mM ADP, and 8 mM MgCl_2_. Reactions were allowed to proceed for 60 min at 30 °C, then the reactions were concentrated in a 100 MWCO spin filter. Concentrated reactions were separated by size exclusion chromatography on a 1 cm x 30 cm Superdex S-200 column equilibrated with anaerobic 100 mM MOPS, 50 mM Tris, 50 mM NaCl (pH 7.3). Less than 500 μL of concentrated reaction was injected, and products were eluted at a rate of 0.5 mL/min. Peaks were monitored by absorbance of 410 nm.

### Nitrogenase turnover assays

Nitrogenase activity was determined by acetylene reduction assay to ethylene as measured in a 9 mL headspace. MoFe-protein and Fe-protein were incubated at varying CRs in reactions with 20 mM sodium dithionite, 5 mM MgCl_2_, 5 mM ATP, 20 mM creatine phosphate, 23 U/mL creatine phosphokinase, buffered in 50 mM Tris-HCl (pH 7.8) with 200 mL NaCl. One mL reactions were incubated at 30 °C for 10 min in an argon atmosphere with 1 mL acetylene gas. Reactions were quenched with 0.25 mL 1 M citric acid and ethylene was quantified by gas chromatography of 40 μL aliquots removed from the head space.

### CryoEM sample preparation and data collection

Solutions of MoFe-protein alone and stabilized complex were prepared in 50 mM Tris-HCl, 200 mM NaCl, 5 mM sodium dithionite (pH 7.8) in an anaerobic chamber. Briefly, 3-4 μL of sample were applied to freshly glow-discharged Quantifoil R1.2/1.3 300 mesh ultrathin carbon grids and blotted for 1-3 seconds with a blot force of 6, at ∼90% humidity using a Vitrobot Mark IV (FEI) in an anaerobic chamber. For aerobic preparation of MoFe-protein grids, the protein solution was prepared in 50 mM Tris-HCl, 200 mM NaCl, 5 mM sodium dithionite (pH 7.8) in an anaerobic chamber. This solution was transferred to a 10 mL Wheaton vial and capped with rubber septa. The Wheaton vial was removed from the anaerobic chamber. Three-four μL of protein solution was removed from the Wheaton vial using a Hamilton gas tight syringe equilibrated with an anaerobic solution of 5 mM sodium dithionite in 50 mM Tris-HCl, 200 mM NaCl (pH 7.8). The protein solution was immediately transferred onto a grid within an aerobic benchtop Vitrobot Mark IV (FEI) and plunge frozen using the blotting conditions described above. Grids were stored in liquid nitrogen until data collection. The MoFe-protein alone and stabilized complex datasets were collected with a 6k x 4k Gatan K3 direct electron detector and Gatan energy filter on a 300 keV Titan Krios in superresolution mode using SerialEM (*35*) at a pixel spacing of 0.65 Å. A total dose of 60 e-/Å^2^ was utilized with a defocus range of −0.8 to −3.0 uM at the Caltech CryoEM Facility.

### CryoEM image processing

Processing of all datasets was performed in cryoSPARC 3 (*36*). Movie frames were aligned and summed using patch motion correction, and the contrast transfer function (CTF) was estimated using the patch CTF estimation job. Templates for automated picking in cryoSPARC were generated by manually picking from a subset of micrographs. For the anaerobic, detergent-containing (dataset 1) and the anaerobic, ultrathin-carbon (dataset 2) MoFe-protein alone datasets 5,724,570 and 4,444,946 particles were picked from 6,777 and 5,130 micrographs, respectively. For the aerobic, ultrathin-carbon (dataset 3) MoFe-protein alone dataset 6,367,415 particles were picked from 8,268 micrographs. For the anaerobic, ultrathin-carbon dataset of the stabilized complex (dataset 4) 2,909,214 particles were picked from 11,669 micrographs. In each case, particles were used for three rounds of reference-free 2D classification and heterogeneous refinement. This yielded final particle subsets of 137,629; 241,057; 219,537; and 36,762 for datasets 1 through 4, respectively. Global CTF refinement was carried out on final subsets of particles, followed by non-uniform refinement (*36*). Values for B-factor sharpening were determined by Mrc2Mtz program and applied to unsharpened maps in cryoSPARC (*37*). Resolutions were estimated by the gold-standard Fourier shell correlation (FSC) curve with a cut-off value of 0.143. Local resolution was calculated using the local resolution estimation job in cryoSPARC.

### Model building and refinement

Initial model fitting was carried out in ChimeraX (*38*) using PDB 3U7Q for the MoFe-protein alone maps or the 1N2C for the stabilized complex. Multiple iterations were carried out of manual model building and ligand fitting in Coot (*39*), and refinement in Refmac5 (*40*) and Phenix (*41*). Data collection, refinement, and validation are presented in Table 1.

### CryoET data collection and processing

Grids of purified proteins were prepared as described above and imaged in a Talos Arctica electron microscope operating at 200 keV with a 6k x 4k Gatan K3 direct electron detector. Datasets were collected at a pixel size of 1.47 Å using a dose-symmetric acquisition scheme from +63° to −63° within SerialEM to a cumulative dose of 120 e-/Å^2^ per tilt series. Frames were aligned, motion- and CTF-corrected in Warp (*42*). Tomograms from resulting image stacks were reconstructed in IMOD (*43*) using weighted back projection after patch track alignment. Missing wedge was corrected in IsoNet (*44*). Particles were visualized in these 3D volumes in ChimeraX.

## Data and materials availability

The single particle cryoEM maps and models have been deposited into the PDB and EMDB for release upon publication.

## Figure preparation and presentation

Structural figures were prepared in ChimeraX and Pymol (*45*). Figures were compiled in Adobe Illustrator.

## Acknowledgments

The foundational contributions of Dr. Thomas Spatzal and Dr. Belinda Wenke to establishing the anaerobic Vitrobot system and conducting the original cryoEM structures of nitrogenase are gratefully acknowledged. We thank Dr. Jens Kaiser, Dr. Songye Chen, Ailiena Maggiolo, Przemyslaw Dutka, and Dr. James Howard for invaluable discussions. The generous support of the Beckman Institute for the Caltech CryoEM Resource Center was essential for the performance of this research.

## Supplemental Tables & Figures

**Supplemental Tables & Figures:**
|  | **MoFe-protein alone in CHAPSO detergent** | **MoFe-protein alone on UC layered grids** | **Aerobic MoFe-protein alone on UC layered grids** | **ADP-AIF<sub>4</sub><sup>-</sup> stabilized MoFe-protein – Fe-protein complex on UC layered grids** |
| --- | --- | --- | --- | --- |
| Collection parameters |  |  |  |  |
| Microscope | Titan Krios | Titan Krios | Titan Krios | Titan Krios |
| Camera | Gatan K3 Summit | Gatan K3 Summit | Gatan K3 Summit | Gatan K3 Summit |
| Magnification | 130,000x | 130,000x | 130,000x | 130,000x |
| Voltage (kV) | 300 | 300 | 300 | 300 |
| Recording mode | counting | counting | counting | counting |
| Frames/Movies | 40 | 40 | 40 | 40 |
| Total Electron dose (e-/Å<sup>2</sup>) | 60 | 60 | 60 | 60 |
| Defocus range (μm) | 0.8 - 3.0 | 0.8 - 3.0 | 0.8 - 3.0 | 0.8 - 3.0 |
| Pixel size (Å) | 0.65 | 0.65 | 0.65 | 0.65 |
| Micrographs collected | 6,777 | 5,130 | 8,268 | 11,669 |
| Micrographs used |  |  |  |  |
| Total extracted particles | 5,724,570 | 4,444,946 | 6,367,415 | 2,909,214 |
| Refined particles | 137,629 | 241,057 | 219,537 | 36,762 |
| Symmetry imposed | C2 | C2 | C2 | C2 |
| Nominal Map Resolution (Å) | 2.04 | 2.26 | 1.92 | 2.12 |
| FSC threshold | 0.143 | 0.143 | 0.143 | 0.143 |
| masked/unmasked | 1.9/2.0 | 2.1/2.2 | 1.8/1.9 | 1.9/2.0 |
| Refinement |  |  |  |  |
| Initial model used | 3U7Q | 3U7Q | 3U7Q | 1N2C |
| Number of atoms |  |  |  |  |
| Protein | 17884 | 16817 | 16986 | 26171 |
| Ligand | ICS:2; CLF:2; HCA:2; FE:2; CPS:4 | ICS:2; CLF:2; HCA:2; FE:2 | ICS:2; CLF:2; 1CL:2; HCA:2; FE:2 | ICS:2; CLF:2; HCA:2; FE:2; MG:2; ADP:4; AIF:4; SF4:2 |
| MapCC (mask/box) | 0.90/0.77 | 0.88/0.78 | 0.83/0.75 | 0.85/0.71 |
| Map sharpening B-factor | 40 | 50 | 30 | 30 |
| R.m.s. deviations |  |  |  |  |
| Bond lengths (Å) | 0.002 | 0.003 | 0.003 | 0.006 |
| Bond angles (°) | 0.53 | 0.522 | 0.566 | 0.756 |
| MolProbity score | 1.28 | 1.34 | 1.58 | 1.96 |
| Clashscore (all atom) | 5.24 | 6.15 | 9.2 | 8.86 |
| Rotamer outliers (%) | 0.75 | 0.7 | 0.81 | 1.81 |
| Ramachandran plot |  |  |  |  |
| Favored (%) | 98.09 | 98.14 | 97.54 | 95.84 |
| Allowed (%) | 1.91 | 1.86 | 2.46 | 4.03 |
| Outliers (%) | 0 | 0 | 0 | 0.13 |

**Supp. Figure 1:**
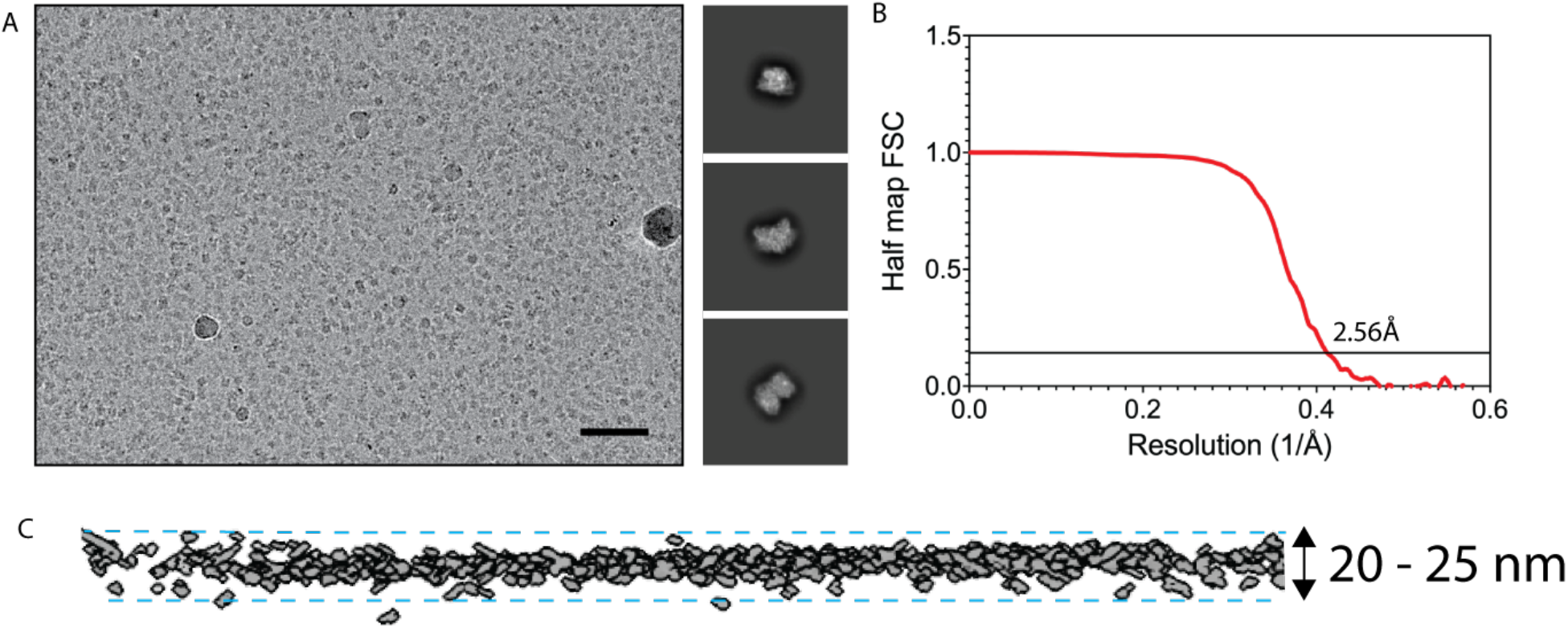
MoFe-protein particles adsorb to the air water interface on holey carbon grids. (a) Left panel, representative micrograph of MoFe-protein particles, scale bar corresponds to 50 nm. Right panel, representative 2D classes of reconstructed volume. (b) Half map FSC curve for reconstructed volume with C1 symmetry. Curve correlates to FSC calculated with a tight mask. (c) Tomographic reconstruction of MoFe-protein particles (gray) in thin ice section. Ice surface is approximated with dashed lines.

**Supp. Figure 2:**
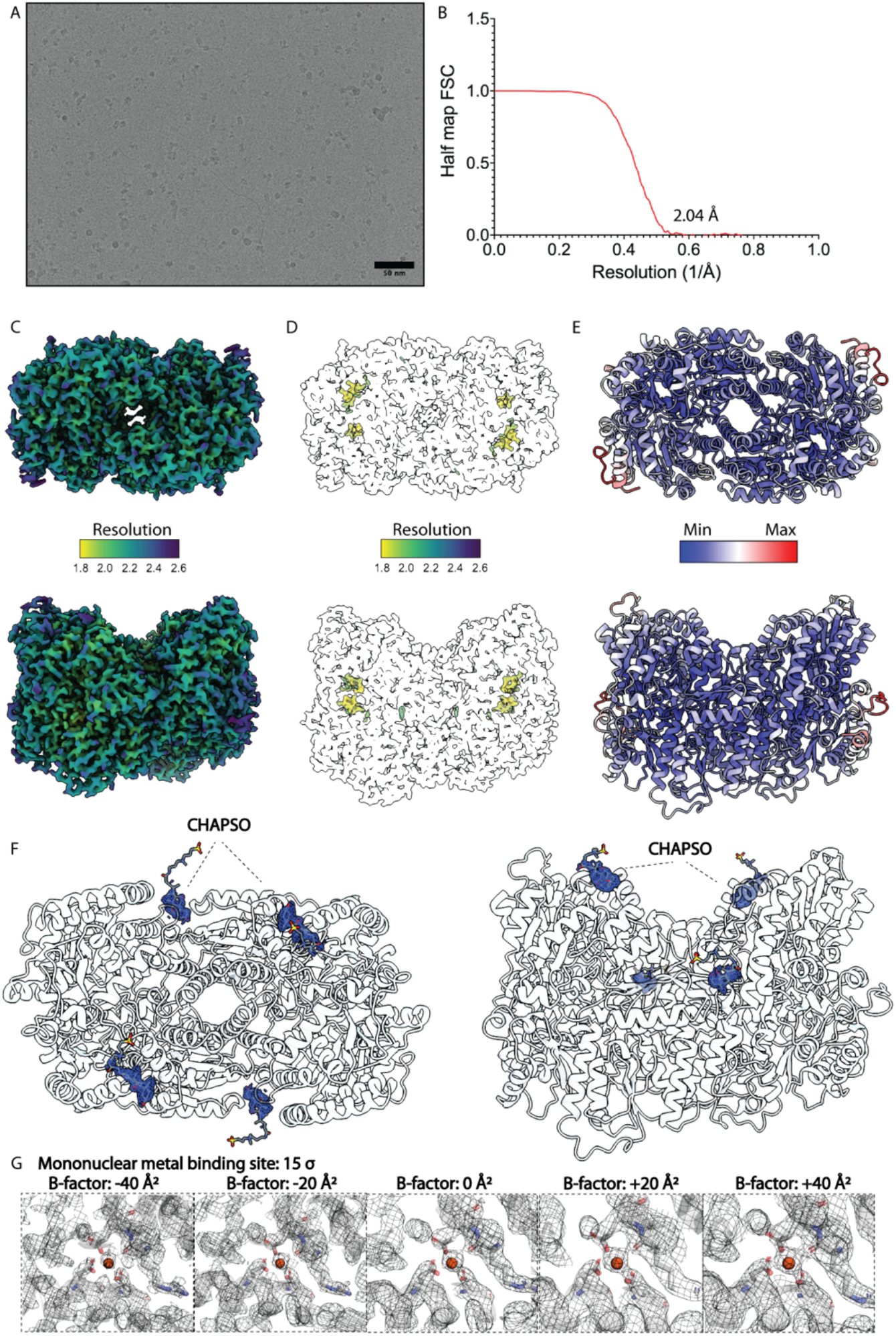
Anaerobic, as-isolated MoFe-protein reconstruction with detergent. (a) Representative micrograph of MoFe-protein particles on holey carbon grids with CHAPSO. Scale bar corresponds to 50 nm (b) Half map FSC curve for reconstructed volume with C2 symmetry imposed. Curve correlates to FSC calculated with a tight mask. (c) CryoEM map color-coded according to local resolution estimates. (d) CryoEM map color-coded according to local resolution estimates at high threshold showing local resolution at metallocluster sites. (e) Model color-coded according to B-factor. (f) ESP for bound CHAPSO molecules. (g) CryoEM ESP for mononuclear metal binding site modeled as Fe(III) with the map sharpened or blurred at indicated B-factors.

**Supp. Figure 3:**
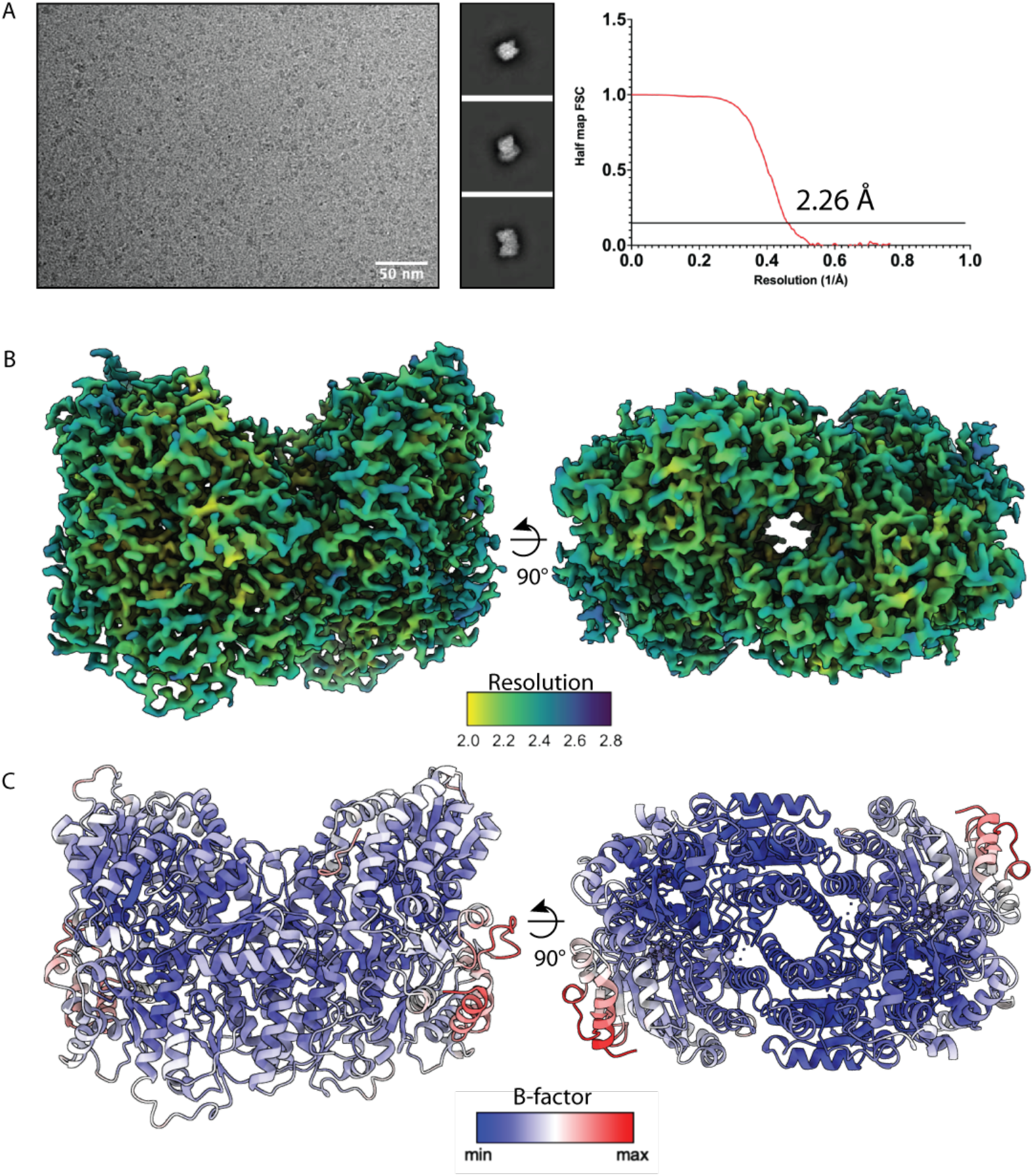
Anaerobic, as-isolated MoFe-protein reconstruction on ultrathin carbon layered grids. (a) Left panel, representative micrograph of MoFe-protein particles on ultrathin carbon layered holey carbon grids. Scale bar corresponds to 50 nm. Middle panel, representative 2D classes. Right panel, half map FSC curve for reconstructed volume with C2 symmetry imposed. Curve correlates to FSC calculated with a tight mask. (b) CryoEM map color-coded according to local resolution estimates. (c) Model color-coded according to B-factor.

**Supp. Figure 4:**
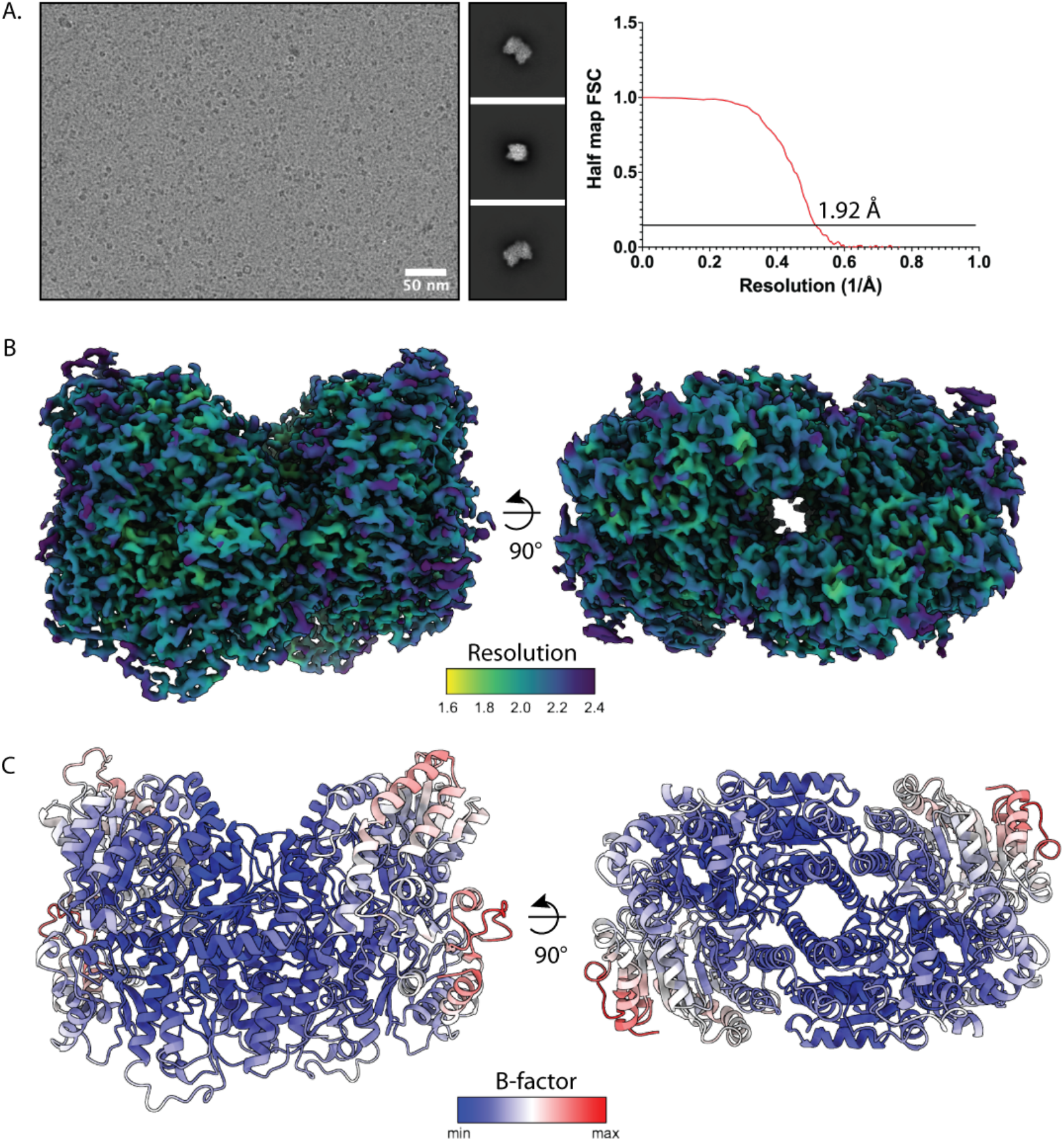
Aerobic as-isolated MoFe-protein reconstruction on ultrathin carbon layered grids. (a) Left panel, representative micrograph of partially oxidized MoFe-protein particles on ultrathin carbon layered holey carbon grids. Scale bar corresponds to 50 nm. Middle panel, representative 2D classes. Right panel, half map FSC curve for reconstructed volume with C2 symmetry imposed. Curve correlates to FSC calculated with a tight mask. (b) CryoEM map color-coded according to local resolution estimates. (c) Model color-coded according to B-factor.

**Supp. Figure 5:**
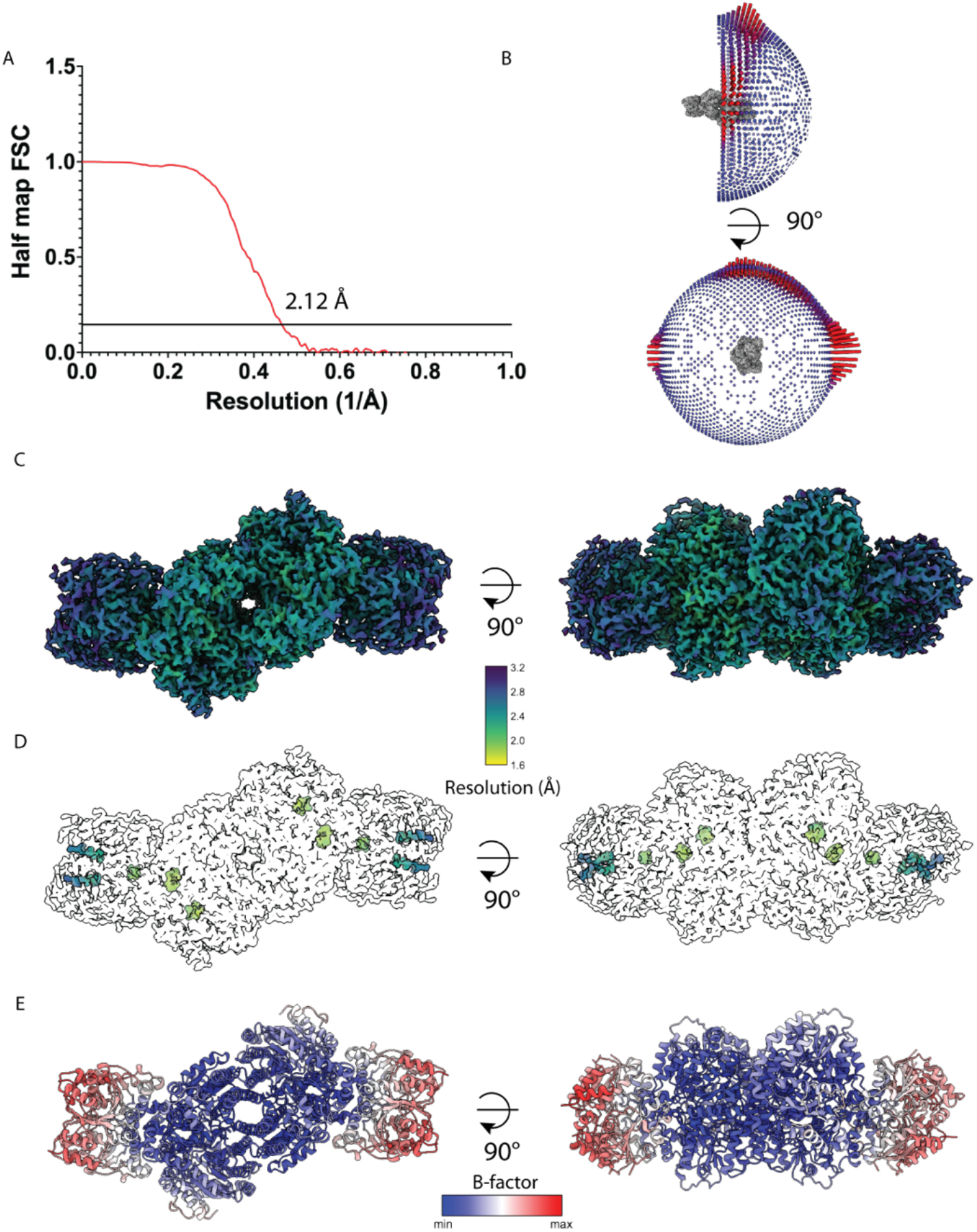
Anaerobic as-isolated ADP-AlF_4_^-^ MoFe-protein – Fe-protein complex on ultrathin carbon layered grids. (a) Half map FSC curve for reconstructed volume with C2 symmetry imposed. Curve correlates to FSC calculated with a tight mask. (b) Euler angle map of particle orientations in reconstructed volume. (c) Map color-coded according to local resolution. (d) CryoEM map color-coded according to local resolution estimates at high threshold showing local resolution at metallocluster sites. (e) Model color-coded according to B-factor.

